# Pout2Prot: an efficient tool to create protein (sub)groups from Percolator output files

**DOI:** 10.1101/2021.08.11.455803

**Authors:** Kay Schallert, Pieter Verschaffelt, Bart Mesuere, Dirk Benndorf, Lennart Martens, Tim Van Den Bossche

## Abstract

In metaproteomics, the study of the collective proteome of microbial communities, the protein inference problem is more challenging than in single-species proteomics. Indeed, a peptide sequence can not only be present in multiple proteins or protein isoforms of the same species, but also in homologous proteins from closely related species. To assign the taxonomy and functions of the microbial species, specialized tools have been developed, such as Prophane. This tool, however, is not directly compatible with post-processing tools such as Percolator. In this manuscript we therefore present *Pout2Prot*, which takes Percolator Output (.pout) files from multiple experiments and creates protein group and protein subgroup output files (.tsv) that can be used directly with Prophane. We investigated different grouping strategies, and compared existing protein grouping tools to develop an advanced protein grouping algorithm that offers a variety of different approaches, allows grouping for multiple files, and uses a weighted spectral count for protein (sub)groups to reflect abundance. *Pout2Prot* is available as a web application at https://pout2prot.ugent.be and is installable via pip as a standalone command line tool and reusable software library. All code is open source under the Apache License 2.0 and is available at https://github.com/compomics/pout2prot.

## Introduction

In metaproteomics, the study of the collective proteome of whole (microbial) ecosystems, it is important to learn about the taxonomy and functions represented in the community. For this purpose, tools such as Unipept^1^ and Prophane^2^ have been made available to specifically perform downstream annotation of metaproteomic data, while other, more generic tools also provide connections to downstream annotation tools^2–4^. These tools, however, work very differently: while Unipept relies on identified peptides without inferring the corresponding proteins (a peptide-centric approach), Prophane uses protein groups as input (a protein-centric approach). Recently, these two tools were compared in the first multi-lab comparison study in metaproteomics (CAMPI)^5^, which indicated that the choice between these approaches is a matter of user preference.

The process of grouping proteins is unfortunately not as straightforward as it might first appear^6–9^. Identified peptide sequences have to be assembled into a list of identified proteins, but when a peptide can be mapped to multiple proteins, this leads to the protein inference problem^9^. In metaproteomics this problem is exacerbated due to the presence of homologous proteins from multiple species in its necessarily large protein databases^10^. Protein grouping is therefore commonly used to generate a more manageable list of identified protein groups that can be used for further downstream analysis. However, different protein grouping algorithms can be chosen, leading to different lists of protein groups from a single set of identified peptides^6^. In the past, many protein grouping methods have been developed, as reviewed in Audain et al^8^, but these typically do not interface well with post-processing tools like Percolator^11^, which are able to increase the number of peptide-to-spectrum matches (PSMs) due to a better separation of true and false matches^12^. Moreover, the common strategy used by these tools is the *Occam’s razor* strategy, which is not always ideal^5^. We here therefore present a new tool, *Pout2Prot*, which provides users with two relevant protein inference options that are tailored towards metaproteomics use cases: *Occam’s razor* and *anti-Occam’s razor*. *Occam’s razor* is based on the principle of maximum parsimony and provides the smallest set of proteins that explains all observed peptides. Here, however, proteins that are not matched by a unique peptide are discarded and their associated taxonomy and functions, which might actually be present in the sample, are lost. This algorithm is for example used in the X!TandemPipeline^13^. On the other hand, *anti-Occam’s razor* is based on the maximal explanatory set of proteins, where any protein that is matched by at least one identified peptide will be included in the reported protein list. This algorithm is used in, for example, MetaProteomeAnalyzer (MPA)^4^. Unfortunately, there is no simple way to determine *a priori* which algorithm will be optimal, as this can differ from sample to sample^5^. These strategies are visually represented in **Figure 1**.

**Figure 1.**
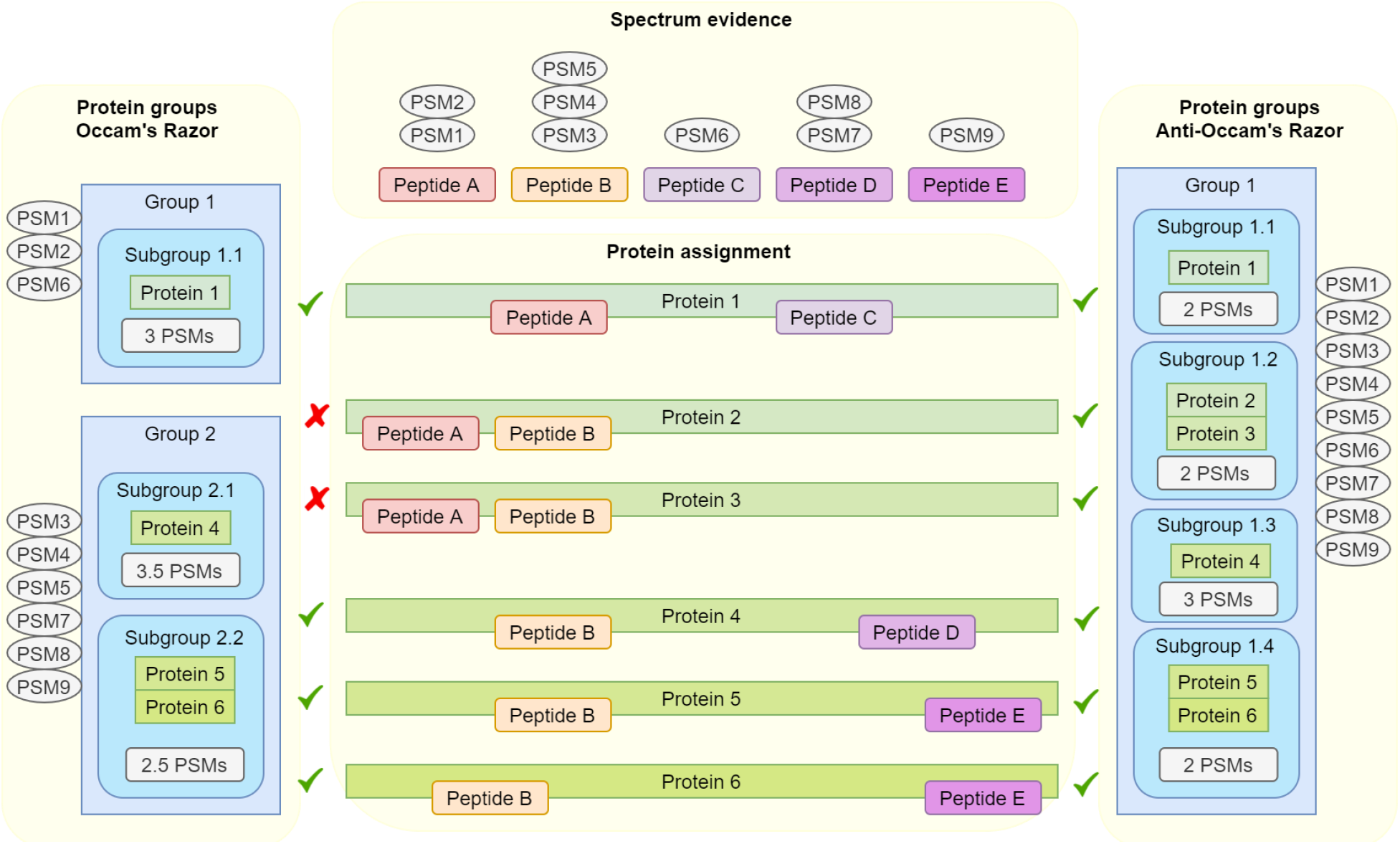
Protein grouping algorithms *Occam’s razor* (left) and *anti-Occam’s razor* (right). Groups can be based on *shared peptide rule* (protein groups) or on *shared peptide set rule* (protein subgroups). This figure also illustrates how PSMs are assigned to protein (sub)groups and shows the weighted PSM count for subgroups. When a PSM is assigned to multiple subgroups, it will be calculated as one divided by the number of subgroups, which can result in fractional PSM counts.

Moreover, as proteins are grouped based on their identified peptides, carefully defined rules are required on when and how to group these proteins. There are two possible approaches here: the first approach consists of grouping all proteins that share one or more identified peptides (i.e., the *shared peptide rule*), while the second approach consists of only grouping proteins that share the same set (or subset) of identified peptides (i.e., the *shared peptide set* rule). These two approaches can also be interpreted as grouping at two different levels: the protein group level (based on the *shared peptide rule*) and the protein subgroup level (based on the *shared peptide set rule*). These two approaches are also visualized in **Figure 1**.

*Pout2Prot* implements all of these approaches: Occam’s razor and anti-Occam’s razor, and this both at the protein group and protein subgroup level. During conceptualization and testing we discovered challenges with the naive description of these algorithms. First, different protein subgroups can have the same peptide and therefore have the same spectrum assigned to them, leading to distorted spectrum counts. Second, when removing proteins using Occam’s razor or when assigning subgroups using anti-Occam’s razor, “undecidable” cases can occur as illustrated in Figure 2. In these undecidable cases, the naive approach might produce inconsistent results when the algorithm is run multiple times.

**Figure 2.**
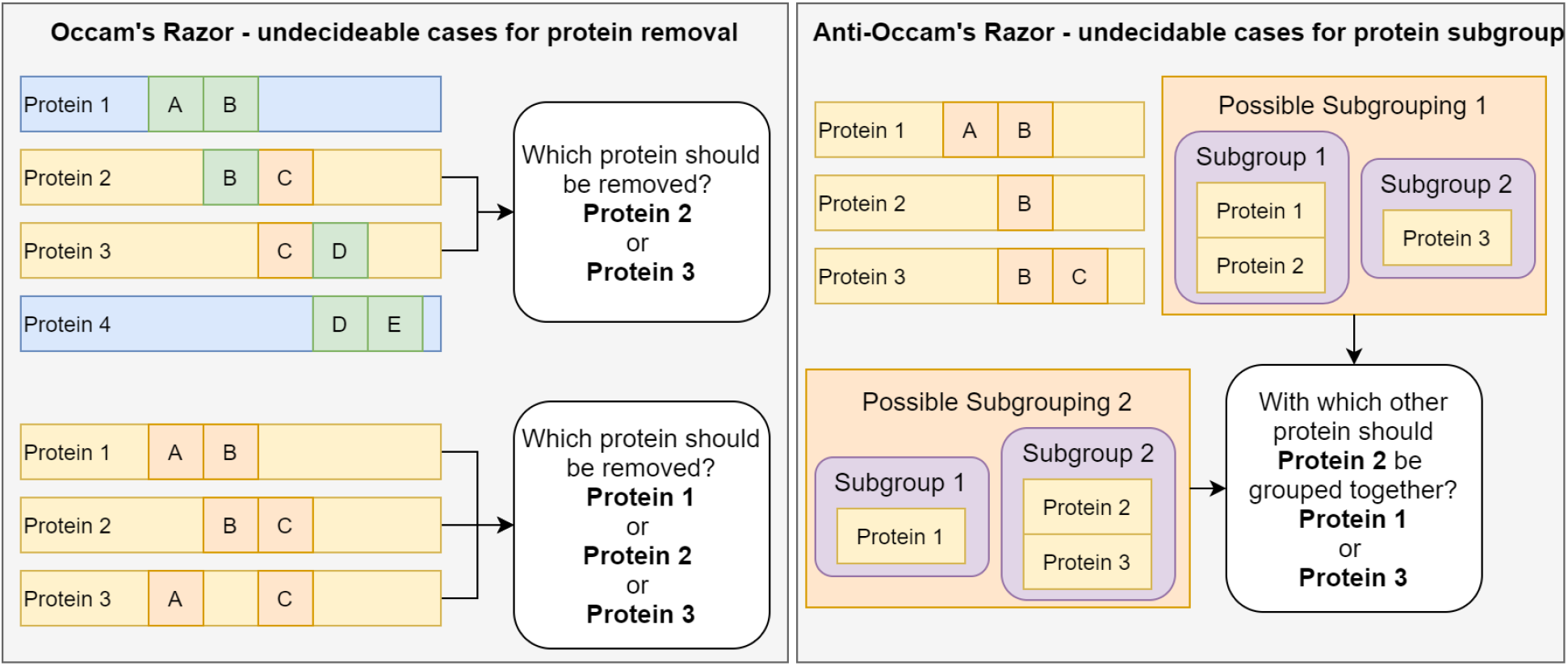
Illustration of undecidable cases. Undecidable cases are situations where peptides and proteins are matched in such a way that the naive interpretation of the algorithm cannot make a clear decision. Specifically, this occurs in Occam’s razor when one of two or more proteins can be removed to explain the remaining peptides (Figure 2, left), and this occurs in anti-Occam’s razor when a protein can be put into a subgroup with two or more other proteins that cannot be subgrouped together (Figure 2, right).

In this manuscript, we describe a new command line tool and web application that can convert .pout files from different experiments into two files containing protein groups and subgroups either as .tsv for direct use with Prophane or as human readable .csv files. Furthermore, we include a file converter that turns Proteome Discoverer output files into the .pout file format. Thus, *Pout2Prot* enables Percolator (or Proteome Discoverer users) to use Prophane for downstream functional and taxonomic analysis.

### Implementation

*Pout2Prot* is implemented in Python and installable as a Python package from PyPI. It can then be invoked from the command line. We also provide a user-friendly and easily accessible web application of our tool (https://pout2prot.ugent.be). The transpiler Transcrypt (https://www.transcrypt.org/) was used to convert our Python package into JavaScript-compatible code and reuse it in our web application. Protein grouping analysis is efficient and can, consequently, be performed entirely on the user’s local machine. Moreover, the web application processes all data locally, so that no data is sent to our servers. This safeguards user data and allows researchers to analyse confidential information more safely.

The detailed implementations of the protein grouping algorithms are visualized in **Supplementary File 1 (Figure 1 and 2)** and consists of four sub-algorithms: the creation of protein groups, the removal of proteins using the rule of maximum parsimony, and a subgroup algorithm each for Occam’s razor and anti-Occam’s razor.

### Evaluation

*Pout2Prot* converts .pout files to protein (sub)group files that can be immediately imported in Prophane for further downstream analysis. This Prophane input file consists of four tab-separated fields: sample category, sample name, protein accessions, and spectrum count. The sample category allows users to divide their experiment in different categories (e.g. “control” and “disease”). If no sample categories are provided, these values will be identical to the sample name, which results in individual quantification by Prophane. The sample name is identical to the name of the .pout file, so each protein (sub)group can be traced back to its origin file. The protein accessions will contain the proteins present in the protein (sub)group, based on the chosen strategy. Finally, the spectrum count contains the weighted spectrum count from all PSMs present in that protein (sub)group, with PSMs present in multiple subgroups counted as fractional values in each subgroup.

#### Qualitative comparison to other tools

To develop a protein grouping algorithm and to truly compare different protein grouping tools, the behaviour of the algorithm must be validated against a set of well defined data, where differences between expected and observed behaviour (i.e. the composition of the groups) can be clearly distinguished. During the development of *Pout2Prot* it quickly became clear that multiple algorithms can solve certain test cases, but fail at others. This also led to the discovery of the undecidable cases outlined in Figure 2. Therefore, we created fourteen test cases (**Supplementary File 2**) that capture all possible pitfalls of protein grouping algorithms, and solved those cases by using both Occam’s razor and anti-Occam’s razor at the protein group and subgroup level. To resolve the issue of undecidability, we propose that no choice should be made at all. For undecidable cases for protein removal (Occam’s razor), no protein should be removed, and for undecidable cases of protein subgroups (anti-Occam’s razor), the protein in question should remain in its own subgroup.

Table 1 shows the result of the comparison between five protein grouping tools: PIA, Fido (integrated into Percolator), MetaProteomeAnalyzer (MPA), X!TandemPipeline, and *Pout2Prot*. To run tests with each tool, appropriate input files that reflect the test cases were created manually and these are all available on the *Pout2Prot* GitHub repository. If a test case did not produce the expected output, it was investigated more closely to ensure this was not the result of differences between, or potential errors in, these input files. For undecidable cases, it was verified that the random choice behaviour could be observed (i.e. multiple analyses, different results). For anti-Occam’s razor subgrouping Cases 3 and 10 a difference in behaviour was observed for PIA and Fido that can be attributed to a different conception of what a protein group is. Specifically, if a protein’s peptide set is a strict subset of another protein’s peptide set, PIA and Fido will not group these two proteins, while MPA and *Pout2Prot* will. Of all the tests that could be run, one resulted in an error: the algorithm for X!TandemPipeline for Case 13. In this case, only one of the six proteins was put into a single group, which leads to a situation, where one of the three peptides was not explained by the resulting groups.

**Table 1.**
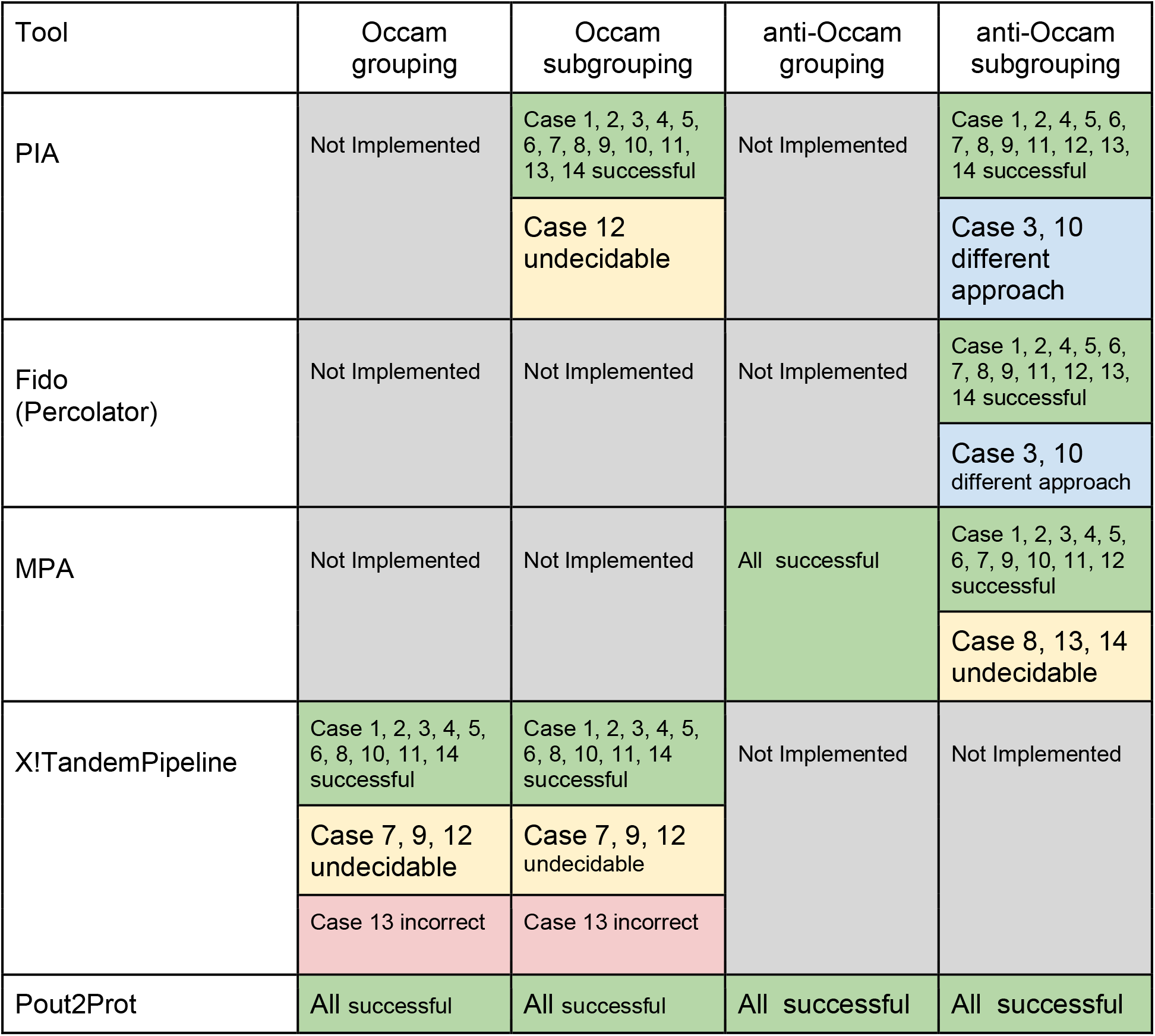
Comparison of the outcome of test cases for five protein grouping tools. The fourteen test cases were run with the PIA, Fido (Percolator), MetaProteomeAnalyzer (MPA), X!TandemPipeline, and *Pout2Prot*. Test cases producing the expected outcome are marked as “successful” (green). Otherwise, these are either categorized as “undecidable” (yellow) if a random choice was made in case of undecidability, “incorrect” (red) if the result cannot be explained logically, and as “different approach” for PIA and Fido, because the anti-Occam protein subgrouping approach used here follows different rules. If a tool does not implement a certain grouping method it is marked as “not implemented” (grey).

While we tried to make a fair comparison, it should be noted that PIA also offers and even recommends another option that falls in between Occam’s razor and anti-Occam’s razor. This method called SpectrumExtractor uses spectrum level information to determine which proteins should be removed or grouped together. Furthermore, Fido offers an option similar to Occam’s razor that operates at the level of the protein database. Percolator and other tools (e.g. Triqler^14^) assign probabilities to proteins instead of making a binary choice for each protein. In contrast, *Pout2Prot* is based on the binary model in which a peptide or protein is either identified or not. This choice is influenced by the fact that a probabilistic approach makes the assignment of taxonomies and functions in metaproteomics very difficult.

#### Performance evaluation

To evaluate the performance of *Pout2Prot*, we tested it on a metaproteomics dataset, derived from the six selected SIHUMIx^14^ datasets used in the Critical Assessment of Metaproteome Investigation (CAMPI) study^5^. Here, we used the X!Tandem^15^ files available on PRIDE^16^ (PXD023217) to (i) convert these files to Percolator Input (.pin) files with tandem2pin, (ii) process the .pin files with Percolator resulting in Percolator Output (.pout) files, and (iii) convert these .pout files to protein (sub)grouping files with *Pout2Prot*, once using Occam’s razor, once using anti-Occam’s razor.

Interestingly, the identification rate (the number of identified spectra divided by the total number of spectra measured) at 1% False Discovery Rate (FDR) increases on average by 7% when using Percolator (**Figure 3a**, blue bars (X!Tandem) vs red bars (Percolator)). It’s important to notice that *Pout2Prot* takes into account the PSM FDR, not the protein FDR. As expected and described before, the semi-supervised machine learning algorithm Percolator is able to increase the number of PSMs due to the better separation of true and false matches^11,12^. More interestingly, we examined the effect of Percolator on the number of protein groups and subgroups. To establish the number of protein (sub)groups before Percolator analysis, we reanalyzed the publicly available raw files of the selected datasets with MPA, also using X!Tandem with identical search settings. Note here that MPA is only able to group proteins according to the anti-Occam’s strategy, so only those numbers were compared in the section below.

**Figure 3.**
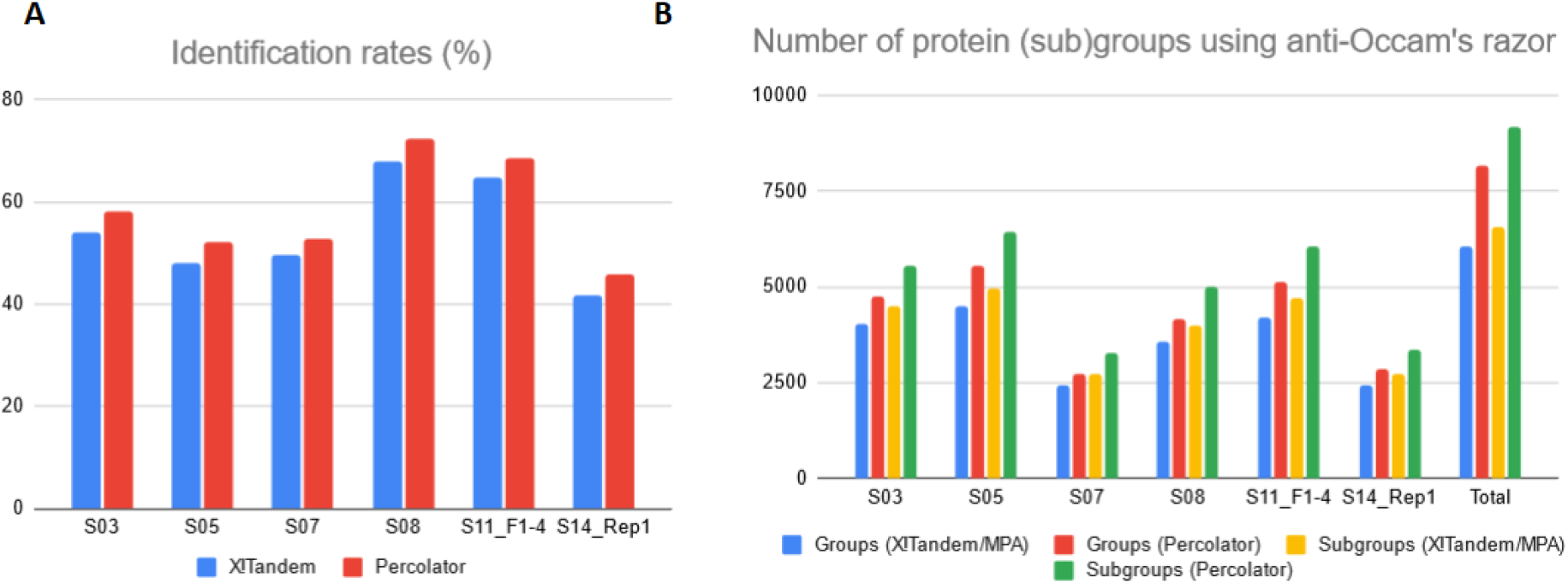
(A) Identification rates per sample for X!Tandem and Percolator analyses. Here, the identification rate was defined as the number of identified spectra divided by the total number of spectra measured. (B) Number of protein (sub)groups compared between X!Tandem and Percolator for the anti-Occam’s razor strategy, and number of protein (sub)groups using Percolator for the Occam’s razor strategy. S03, S05, S07, S08, S11_F1-4, and S14_Rep1 refer to the six SIHUMIx samples.

In **Figure 3b**, we observe that after Percolator analysis, the number of protein groups per sample increased by 18.5% on average (blue vs red bars) and the number of protein subgroups per sample increased by 25.3% on average (yellow vs green bars). The total number of groups and subgroups across all samples increased more drastically (by 34.7% and 39.9%, respectively) in comparison to the averages per sample. All raw data is available in **Supplementary File 3 (Table 1 and 2)**.

Furthermore, we also investigated the effect on the number of protein (sub)groups of combining different fractions at different places in the workflow. We combined (i) the Mascot Generic Format (.mgf) files before the X!Tandem search, (ii) before the Percolator search, and (iii) before *Pout2Prot* protein inference. Since the range for the number of protein (sub)groups constitute 2-3% difference, the point in the workflow where the different files are combined, is of minimal impact (**Supplementary File 3, Table 3**).. For completeness, an example result file for taxonomic and functional analysis after processing of *Pout2Prot* output in Prophane can be found in **Supplementary File 3, Figure 1 and 2**). In addition, the time for a *Pout2Prot* analysis (Occam’s razor) for the complete SIHUMIx experiment via the web service was less than 5 seconds.

## Conclusion

*Pout2Prot* enables the conversion of Percolator output (.pout) files to protein group and protein subgroup files, based on either the Occam’s razor or anti-Occam’s razor strategy, and therefore closes an important gap in the bioinformatic workflow of metaproteomics data analysis. Moreover, *Pout2Prot* also allows the user to create protein (sub)groups across experiments. The output of *Pout2Prot* can be imported directly into Prophane, which in turn allows users to perform downstream taxonomic and functional analysis of metaproteomics samples.

## Supporting Information

The following supporting information is available free of charge at ACS website http://pubs.acs.org.

Supplementary File 1: Algorithmic description

Supplementary File 2: Test cases

Supplementary File 3: Additional results

## Acknowledgements

This work has benefited from collaborations facilitated by the Metaproteomics Initiative (https://metaproteomics.org/) whose goals are to promote, improve and standardize metaproteomics. TVDB, PV, LM and BM would like to acknowledge the Research Foundation - Flanders (FWO) [grants 1S90918N, 1164420N, G042518N and 12I5220N]. LM also acknowledges support from the European Union’s Horizon 2020 Programme under Grant Agreement 823839 [H2020-INFRAIA-2018-1]. KS and DB would like to acknowledge the German Federal Ministry of Education and Research (BMBF) of the project ‘MetaProteomanalyzer Service’ within the German Network for Bioinformatics Infrastructure (de.NBI) [031L103]. The authors declare no conflict of interest.

