## Supplementary File 1 for "Pout2Prot: an efficient tool to create protein (sub)groups from Percolator output files"

**Supplementary File 1: Algorithmic description**

This Supplementary File explains the protein grouping algorithms implemented in Pout2Prot. The overall workflow starts by creating two mappings (Python dictionaries): the protein identifiers are mapped to the set of peptide sequences (for this protein), and the peptides sequence is mapped to the protein identifiers (all proteins that contain this peptide). Then, one of two separate paths are chosen whether or not Occam's razor is selected.
For anti-Occam's razor, the general grouping algorithm and subsequently the special subgrouping algorithm for anti-Occam's razor are executed, to create protein groups and subgroups respectively. For Occam's razor the same, general grouping algorithm is executed first. This allows the Occam Filter algorithm to run much more efficiently because it can be applied to a single group one at a time. The Occam Filter will remove proteins from the two mappings created at the beginning. The second general grouping of the Occam's razor path then produces the correct groups and the subgrouping algorithms the subgroups.

The grouping algorithm works identical in all three occurrences in the workflow and uses a recursive call to identify a subgraph in the network of protein and peptide nodes by continuously expanding until no more proteins and peptides are found that can be added.

The subgrouping for Occam's razor is very straightforward and groups proteins that share the exact same set of peptides. The subgrouping for anti-Occam's razor is more complex since the undecidable cases have to be detected and dealt with. In this algorithm every protein in a specific protein group is checked against every other protein in the group if they can be grouped because one has a strict subset of peptides of the other (i.e protein 1 has peptides A, B, and C and protein 2 has peptides A and B, therefore protein 2 has a strict peptides subset of protein 1). The two proteins are then grouped into the same subgroup if there is a full agreement with which other proteins they can be grouped together (see protein 4 and 5 from **Supplementary File 1, Figure 1, Subgrouping anti-Occam**)

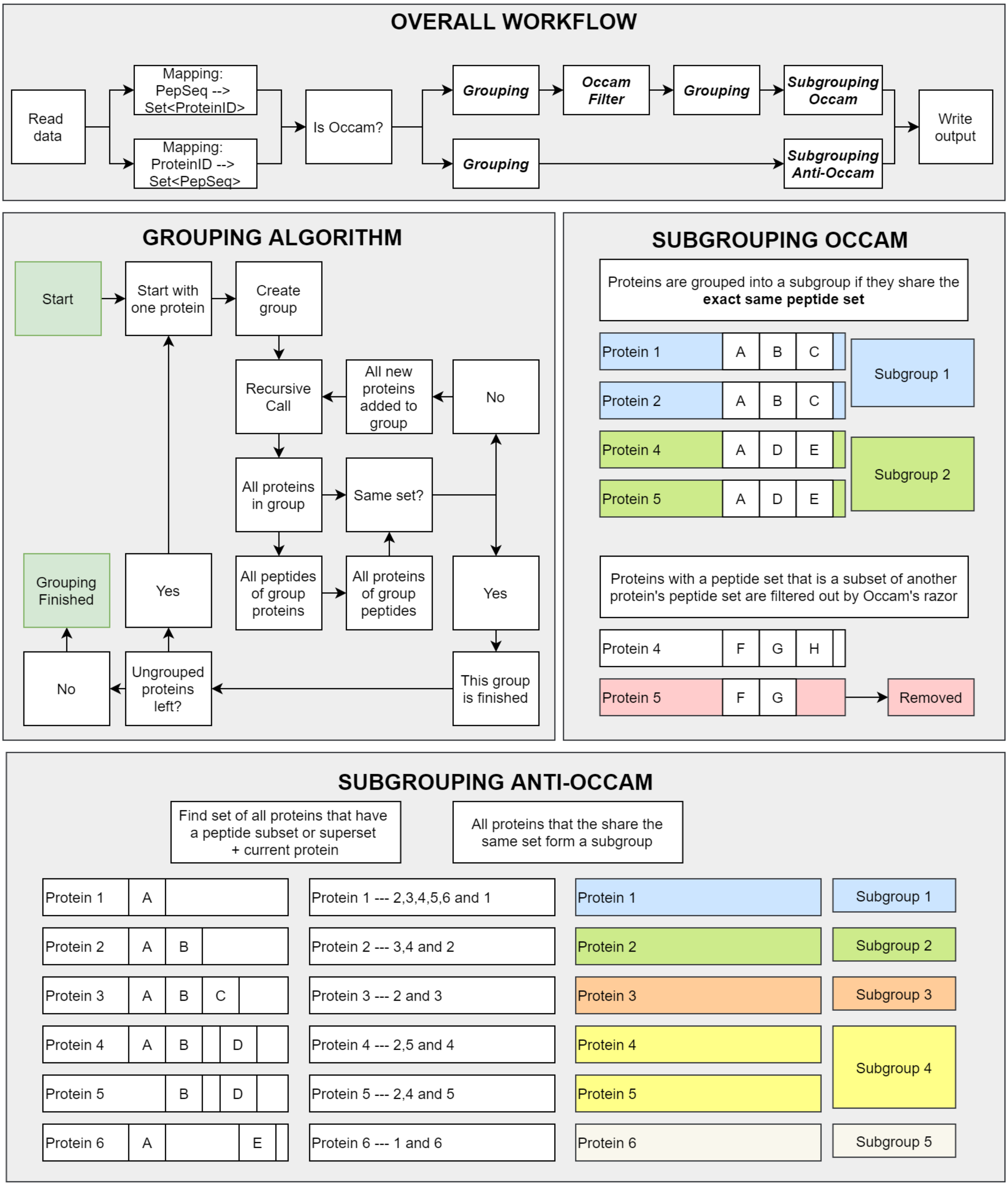


**Supplementary File 1, Figure 1: Algorithmic workflow of Pout2Prot.** The general workflow is visually represented in Overall Workflow (top). The (sub)grouping building blocks of this workflow are visually represented as well: the general grouping algorithm (middle, left), the subgrouping algorithm of Occam's razor (middle, right), and the subgrouping algorithm of anti-Occam's razor (bottom).

The Occam Filter algorithm is the most complex of the algorithms outlined here because it also has to deal with undecidable cases. The algorithm works on a single (anti-Occam) protein group because this includes all proteins that can possibly be connected based on the peptide evidence. The simple and common situations are dealt with in steps 1 - 5. First, duplicate peptides (with the exact same peptide set) are detected and considered as a single protein for the rest of the calculation. Then proteins that have a strict subset of peptides are detected and removed (i.e protein 1 has peptides A, B and C and protein 2 has peptides A and B, therefore protein 2 has a strict peptide subset of protein 1). Proteins that have a unique peptide are detected next, after which it is determined if these proteins with unique peptides already explain all the peptide evidence. If that's the case, all other proteins are removed and the next protein group can be processed.

If the algorithm has to continue, it attempts to create the smallest possible solution for the remaining peptides (peptides that are not explained by proteins with unique peptides). The smallest possible solution is the smallest number of proteins (i.e. the *solution_size*) that explains all remaining peptides. If a single solution is found for a given solution_size, the best solution is found, and unused proteins can be removed. If multiple solutions are available for a given solution_size, the solutions are scored based on the proteins they contain. We consider the score of a protein to be the number of solutions that it is contained in and the score of the solution to be the sum of protein scores. The highest scoring solution is chosen and unused proteins are removed. If the highest score is given to multiple solutions (or there are no solutions for a given solution_size), then the solution_size is increased and the algorithm is repeated until a solution is found. If, finally, no solution is found, no proteins are removed.

The principle behind the choice of this scoring scheme is that if a protein is occurring in more solutions, then it is more important to explain the remaining peptides. In truly undecidable cases all solutions will have the same score, thus, the undecidability is detected. This particular implementation is one of many possibilities to achieve this and we do not claim to have found the single true solution to this problem.

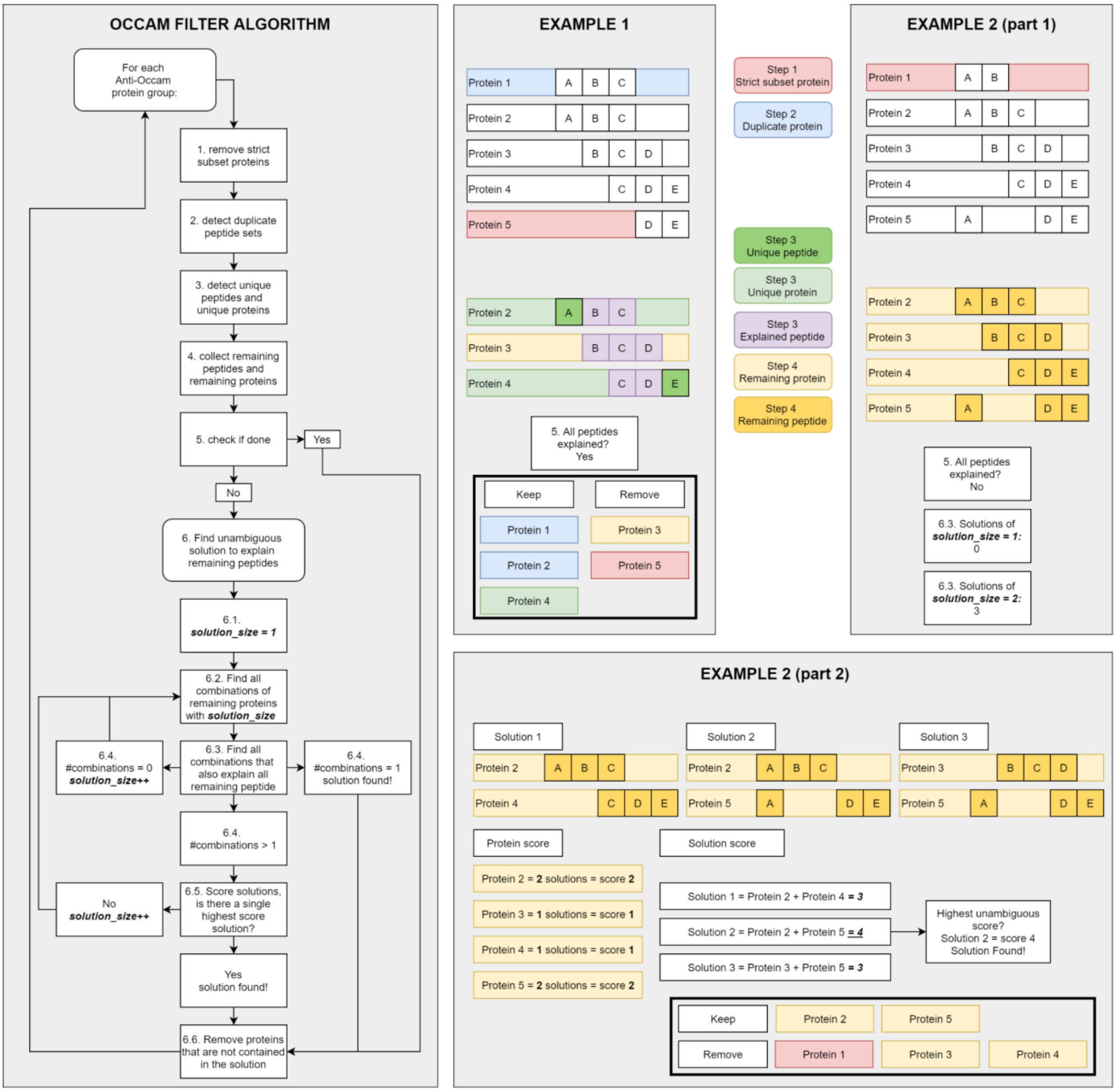


**Supplementary File 1, Figure 2:** The Occam Filter algorithm visually represented (left), including two examples (right).
