## Supplementary File 2 for "Pout2Prot: an efficient tool to create protein (sub)groups from Percolator output files"

**Supplementary File 2: Test cases**

This file visually explains the results of the fourteen test cases used for comparison to other protein grouping tools. The files that were input into each tool representing the test cases are found in data/manuscript/tool-comparison on GitHub. A detailed description of the test cases are given in the figure legend.

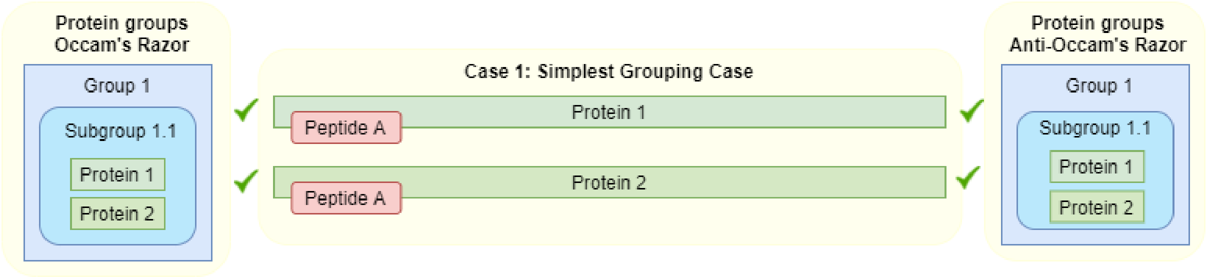


**Supplementary File 2, Figure 1:** Case 1. This is the simplest grouping case: two proteins share the same peptide. Group and subgroup are identical, the rule of parsimony does not apply.

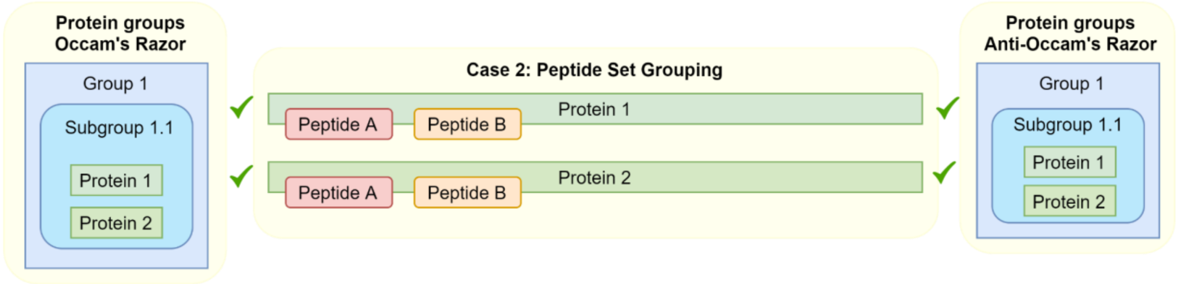


**Supplementary File 2, Figure 2:** Case 2. This grouping case is similar to Case 1, but both proteins share 2 peptides. Group and subgroup are identical, the rule of parsimony does not apply.

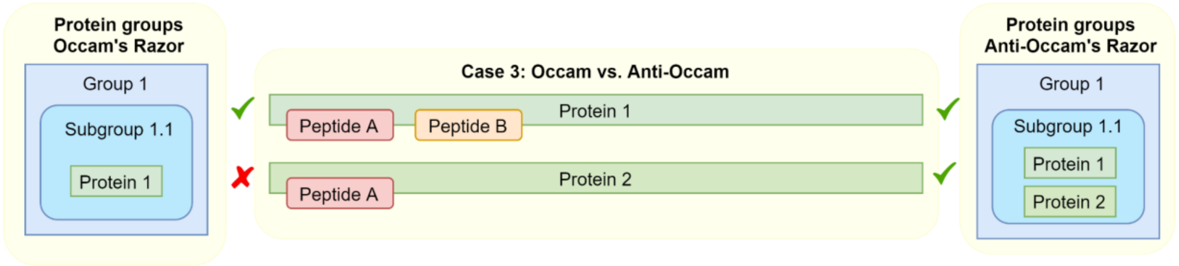


**Supplementary File 2, Figure 3:** Case 3. This grouping case is the simplest application of the parsimony principle: protein 2 must be removed because it does only explain one peptide instead of two that are explained by protein 1. This means there is a difference between the anti-Occam's razor and the Occam's razor approach.

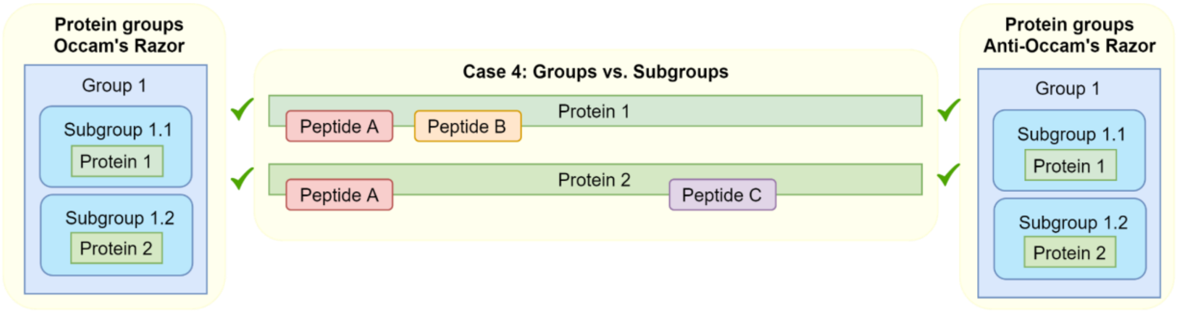


**Supplementary File 2, Figure 4:** Case 4. This grouping case is the simplest application of the group/subgroup distinction. A group only requires that proteins share at least one peptide. A subgroup requires that proteins share an entire peptide set (and have no unique peptides). Because protein 1 and protein 2 share peptide A, they should be grouped into the same group, but because they possess a unique peptide each (B and C), they should be grouped into different subgroups.

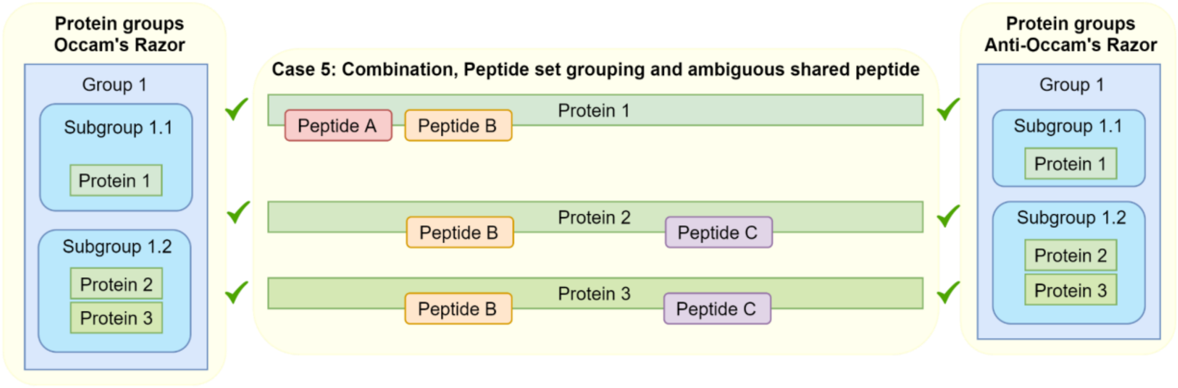


**Supplementary File 2, Figure 5:** Case 5. This grouping case shows how a subgroup is formed within a larger group. All 3 proteins are connected by peptide B and are therefore grouped together. Protein 1 has a unique peptide and forms its own subgroup. Proteins 2 and 3 share the exact same peptide set and therefore form a subgroup.

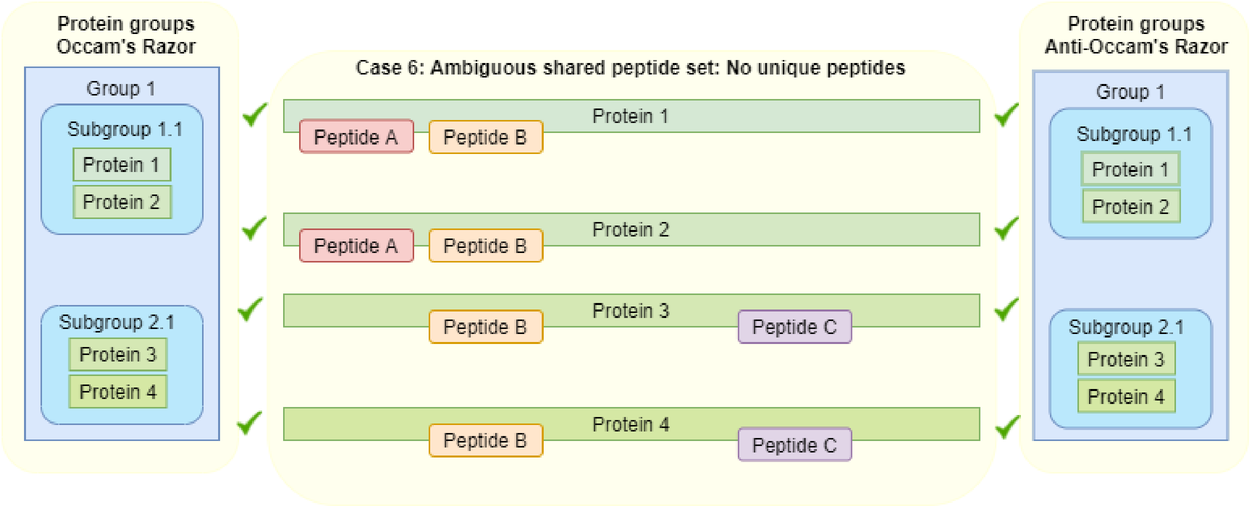


**Supplementary File 2, Figure 6:** Case 6. This grouping case is similar to case 5 but it includes an additional protein that has the same peptide set (A and B) as protein 1 from case 5. This case is considered here, because it contains no unique peptides at all.

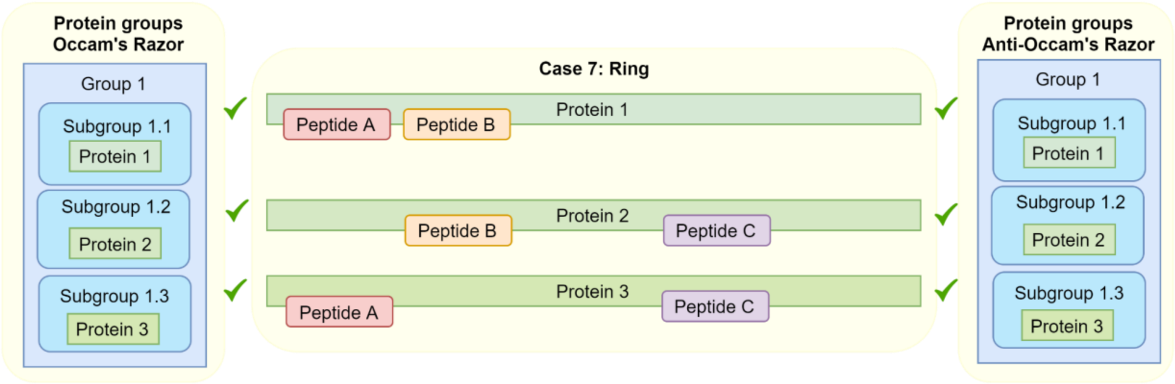


**Supplementary File 2, Figure 7:** Case 7. This grouping case is a special case that can easily break grouping algorithms: a ring. As can be seen, no protein has a unique peptide, no protein has a peptide set that is a subset of another protein's peptide set. The only reasonable course to take here is to put every protein in its own subgroup. Occam-based grouping algorithms can break here by removing one of the proteins through the wrong application of the rule of parsimony (which protein depends on the order of processing the proteins).

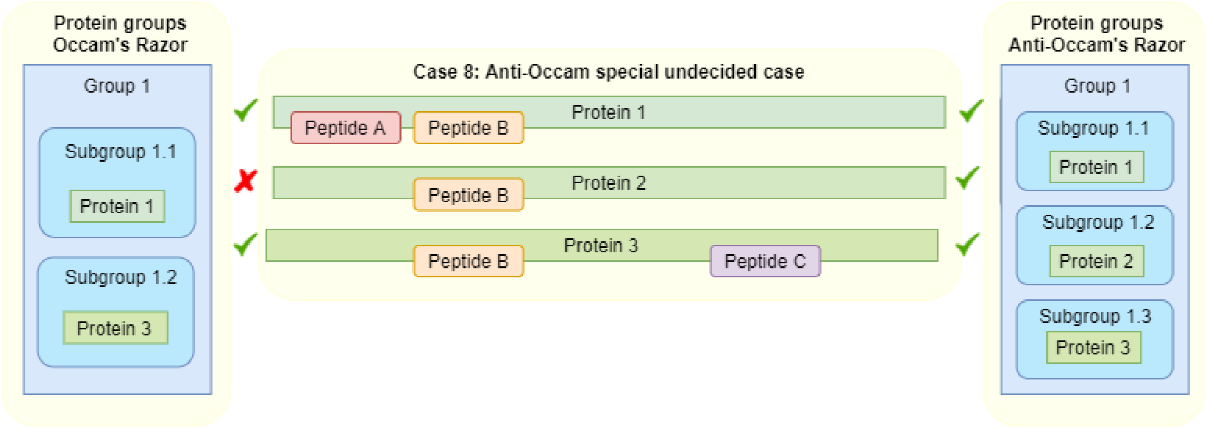


**Supplementary File 2, Figure 8:** Case 8. This grouping case is a special case that can easily break anti-Occam grouping algorithms. As can be seen, the peptide set of protein 2 (just protein B) is a subset of both, the peptide sets of protein 1 and 3. The only reasonable approach here (anti-Occam) is to put protein 2 in its own subgroup. anti-Occam algorithms can break here by putting protein 2 in a subgroup together with either protein 1 or 3 depending on the order of processing.

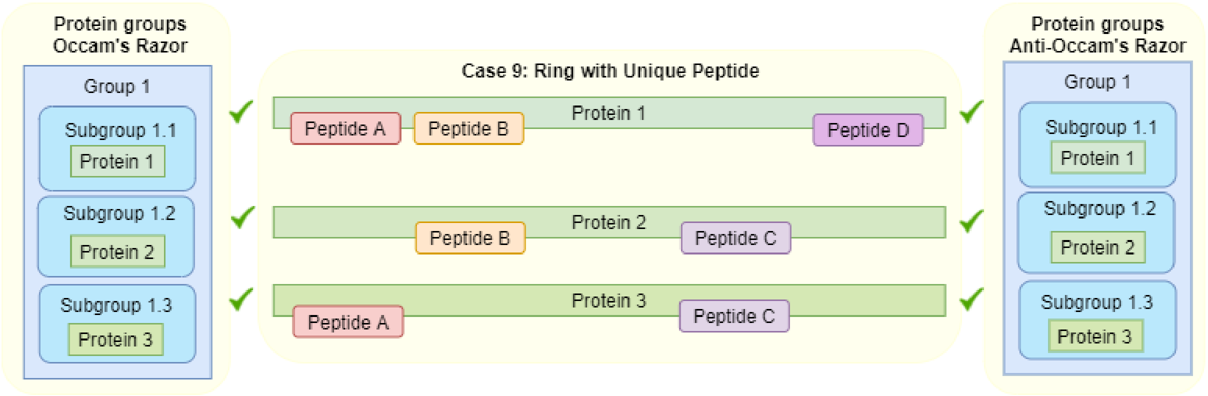


**Supplementary File 2, Figure 9:** Case 9. This grouping case is the ring case (case 7) with an additional unique peptide (peptide D) for protein 1. The inclusion of this peptide can disrupt some implementations that detect rings.

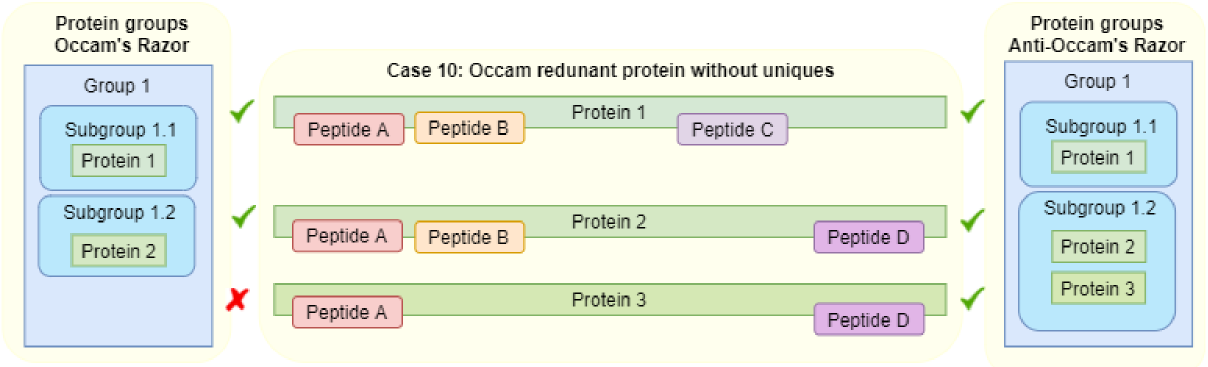


**Supplementary File 2, Figure 10:** Case 10. This grouping case is a more complex case, where grouping algorithms can easily fail, because of the order in which certain steps to create groups are done by the algorithm.

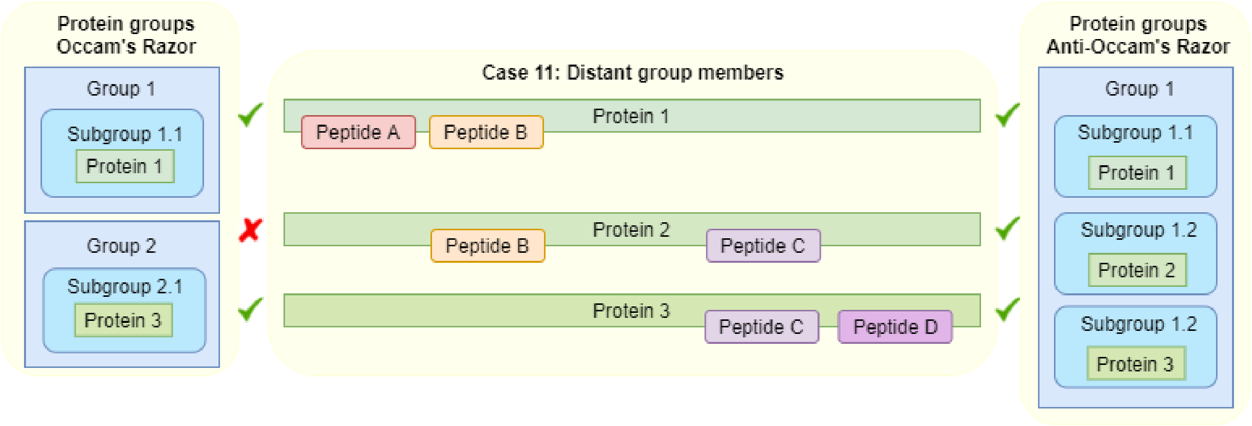


**Supplementary File 2, Figure 11:** Case 11. This grouping case deals with distant group members, meaning that certain proteins in the group don't share a single peptide, in this case protein 1 and 3. Applying the rule of parsimony will separate the group in this specific case. In the anti-Occam case, protein 2 will remain in a separate subgroup.

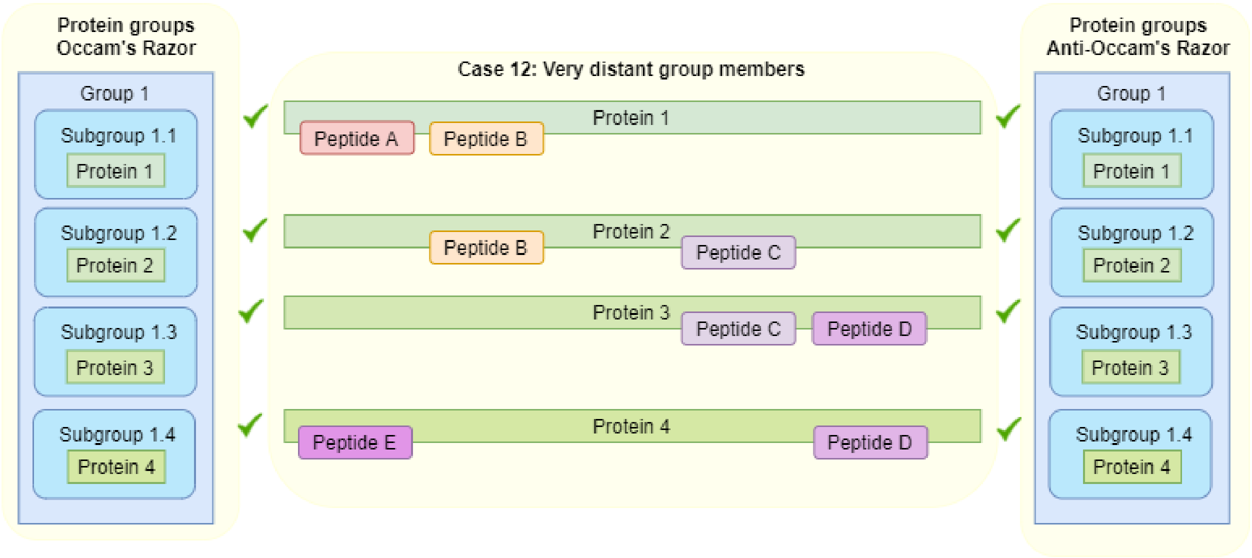


**Supplementary File 2, Figure 12:** Case 12. This grouping case is very similar to case 11, adding one more protein to the chain. Unlike in case 11, the rule of parsimony cannot be applied, because it is unclear which protein should be removed (protein 2 or 3). Incorrect implementations of the parsimony rule will remove either one (randomly chosen, 2 or 3), which will lead to inconsistent grouping if the experiment is repeated. The correct solution is the same grouping for Occam and anti-Occam.

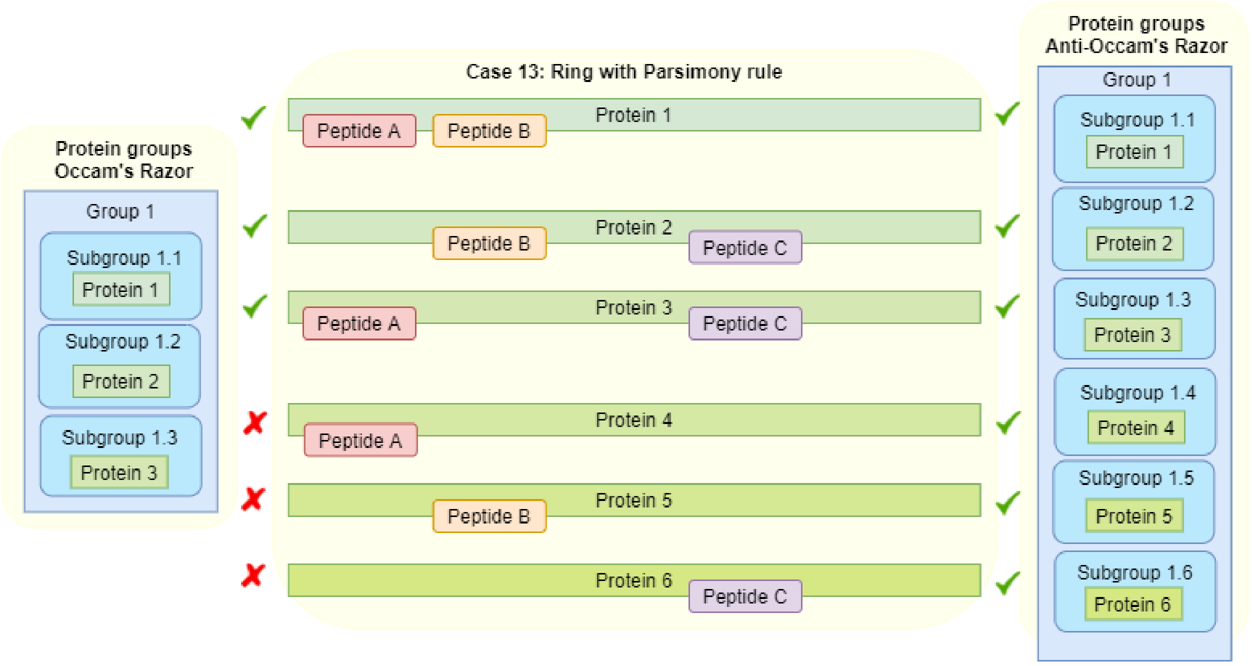


**Supplementary File 2, Figure 13:** Case 14. This is a more complex grouping case; a ring (case 7), but additional redundant proteins are identified (4, 5 and 6). Applying the rule of parsimony should reduce the problem to case 7, but for anti-Occam this will lead to the situation of case 8. Protein 4 could be grouped either with protein 1 or 3, protein 5 could be grouped either with protein 1 or 2 and protein 6 could be grouped either with protein 2 or 3. Since all 6 proteins cannot clearly be assigned to a subgroup, they should remain separate.

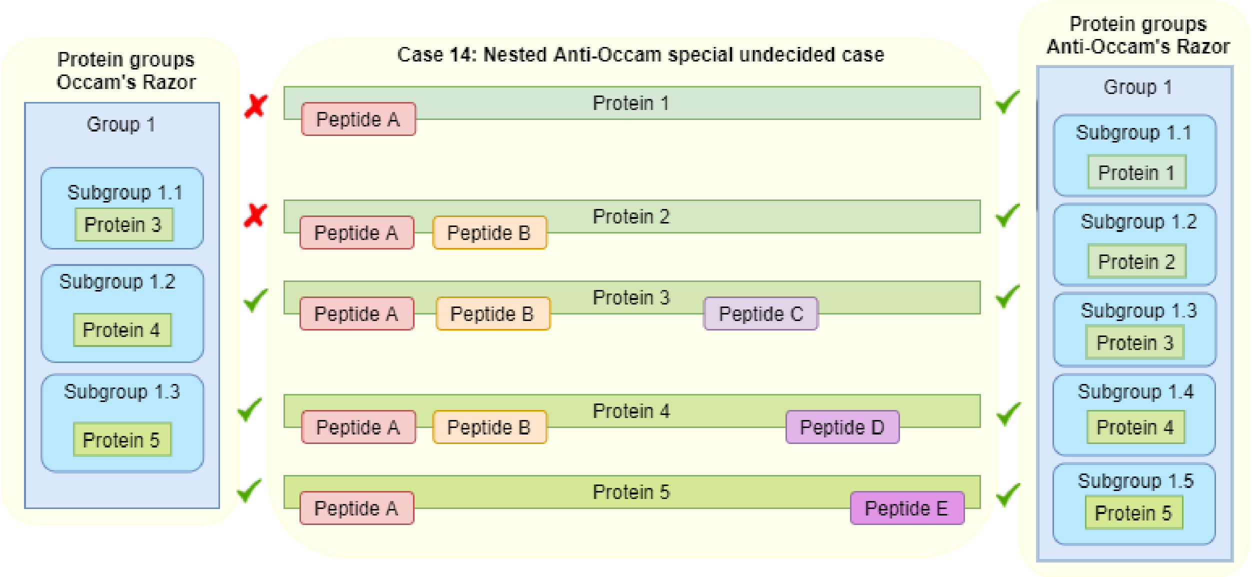


**Supplementary File 2, Figure 14:** Case 14. This grouping case is another more complex example, a nested version of case 8. Like in case 8, protein 1 could be assigned into a subgroup together with either protein 2 or protein 5 - no clear decision can be made. This is true for protein 2 as well, which could be grouped together with either protein 3 or protein 4. In the anti-Occam approach, no protein can be assigned to a clear subgroup. This can break certain algorithms that do not properly account for case 8. The Occam’s razor approach can easily group proteins after removing proteins 1 and 2.
