## Supplementary File 3 for "Pout2Prot: an efficient tool to create protein (sub)groups from Percolator output files"

### Supplementary File 3: Additional results

#### **Supplementary Table 1:** identification rates of X!Tandem and Percolator

| sample | Spectra | X!Tandem PSM count | X!Tandem ID Rate (%) | X!Tandem Pep count | Percolator PSM count | Percolator ID Rate (%) | Percolator Pep count |
|---|---|---|---|---|---|---|---|
| S03 | 157489 | 85300 | 54 | 28591 | 91848 | 58 | 31078 |
| S05 | 240740 | 115893 | 48 | 32546 | 125366 | 52 | 35831 |
| S07 | 46847 | 23307 | 50 | 16215 | 24685 | 53 | 17241 |
| S08 | 81654 | 55458 | 68 | 21535 | 59214 | 73 | 23221 |
| S11_Fraction1 | 43447 | 28808 | 66 | 16012 | 30652 | 71 | 71 |
| S11_Fraction2 | 45613 | 30306 | 66 | 16784 | 32041 | 70 | 70 |
| S11_Fraction3 | 46941 | 31392 | 67 | 17189 | 32939 | 70 | 70 |
| S11_Fraction4 | 37175 | 21585 | 58 | 9800 | 22943 | 62 | 62 |
| S11_F1-4_presearch | 173176 | 112223 | 65 | 39195 | 118687 | 69 | 41803 |
| S11_F1-4_prepercolator | 173176 | | | | 118614 | 68 | 68 |
| S11_F1-4_postpercolator | 173176 | | | | 118575 | 68 | 68 |
| S14_Rep1 | 71265 | 29743 | 42 | 14986 | 32757 | 46 | 16604 |

#### **Supplementary Table 2:** Number of protein (sub)groups

| Sample | anti-Occam Groups (X!Tandem/MPA) | anti-Occam Groups (Percolator) | anti-Occam Subgroups (X!Tandem/MPA) | anti-Occam Subgroups (Percolator) | Occam Groups (Percolator) | Occam Subgroups (Percolator) |
|---|---|---|---|---|---|---|
| S03 | 4044 | 4735 | 4503 | 5552 | 8178 | 9109 |
| S05 | 4483 | 5559 | 4950 | 6429 | 4735 | 5525 |
| S07 | 2420 | 2731 | 2724 | 3289 | 5559 | 6395 |
| S08 | 3550 | 4177 | 3985 | 4987 | 2731 | 3272 |
| S11_F1-4 | 4223 | 5132 | 4692 | 6050 | 4177 | 4945 |
| S14_Rep1 | 2429 | 2860 | 2720 | 3371 | 5132 | 5977 |
| Total | 6070 | 8178 | 6575 | 9197 | 2860 | 3358 |

**Supplementary Table 3:** Comparison of number of protein (sub)groups of fractionation experiment.

| Sample | Occam Groups | Occam Subgroups | anti-Occam Groups | anti-Occam Subgroups |
|---|---|---|---|---|
| S11_F1-4_presearch | 5244 | 5808 | 5244 | 5900 |
| S11_F1-4_prepercolator | 5240 | 5791 | 5227 | 5883 |
| S11_F1-4_postpercolator | 5227 | 5804 | 5240 | 5898 |
| Total | 5387 | 5951 | 5387 | 6045 |

**Supplementary Figure 1:** example of taxonomic distribution Krona plot from Prophane (Occam Subgroups SIHUMIx, S05)

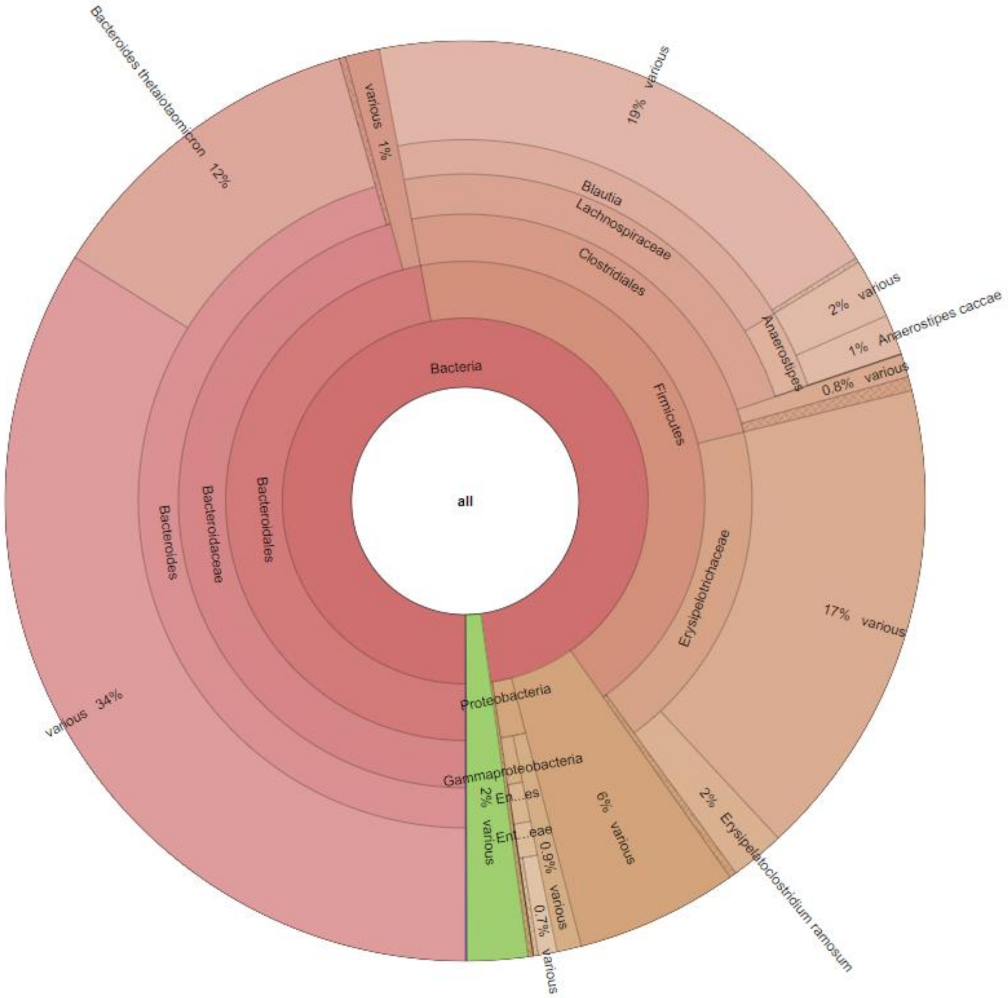

**Supplementary Figure 2:** example of functional distribution Krona plot from Prophane (Occam Subgroups SIHUMIx, S05)

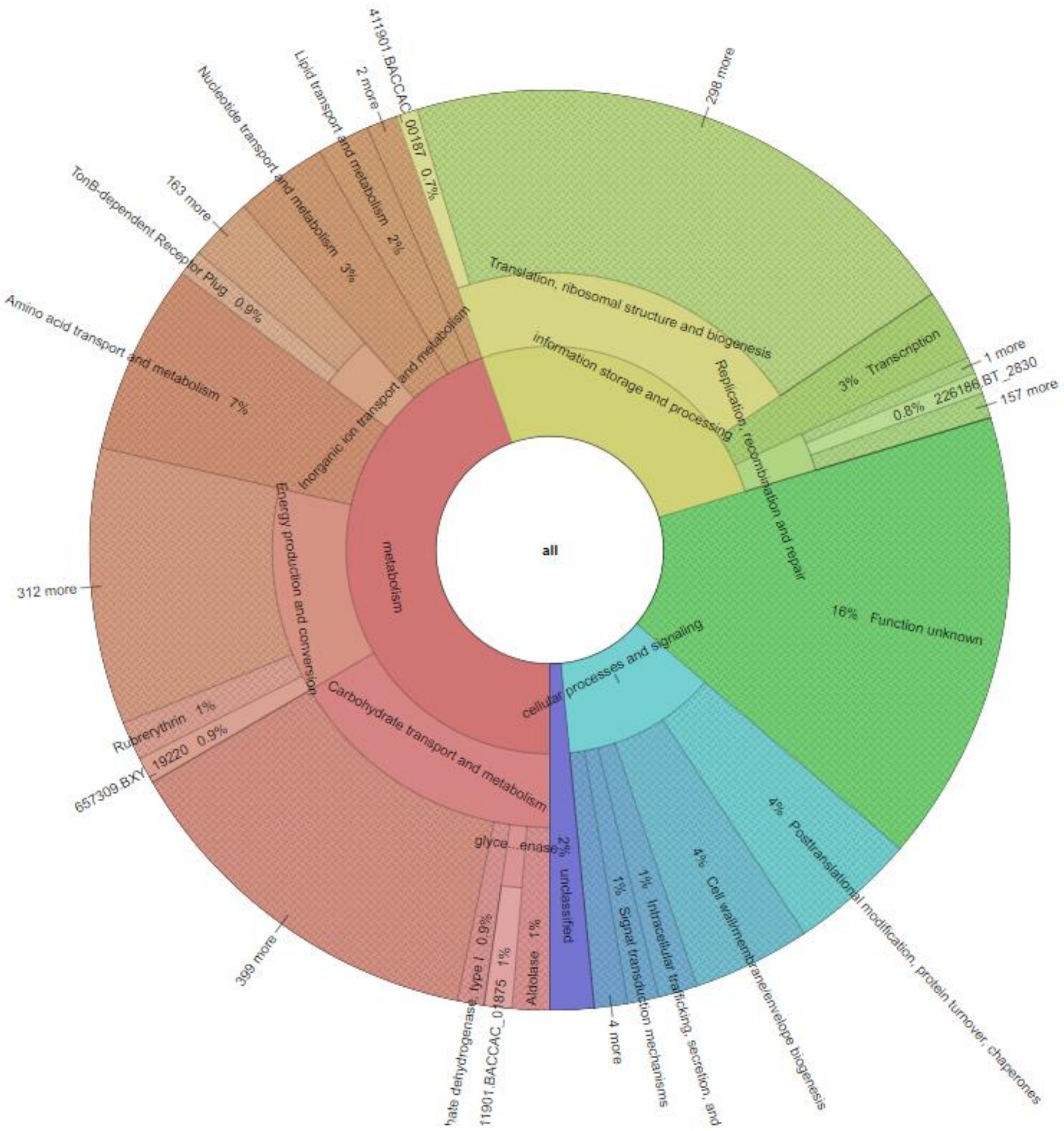
